# Identification of Bone Morphogenic Protein (BMP15) gene at exons 1 and 2 in Indigenous, Mozaffornagri, and Dorper ram breeds of Bangladesh and its phylogenetic relationship with other breeds

**DOI:** 10.1101/2025.04.10.648142

**Authors:** Muhammad Rakibul Hasan, Mohammad Musharraf Uddin Bhuiyan, Nasrin Sultana Juyena, Farida Yeasmin Bari

**Affiliations:** Bangladesh Agricultural University

## Abstract

Sheep play a vital role in alleviating poverty and providing protein for the people of Bangladesh and they are popular for their mutton production. This study aimed to explore the polymorphisms in the BMP15 gene among various Bangladeshi ram breeds for devising genetic improvement strategies to boost mutton output and help reduce poverty. We collected blood samples from 25 rams, including the Indigenous, Mozaffarnagri, and Dorper breeds, all raised in a traditional farming system. After extracting DNA, we amplified the BMP15 gene at Exon-1 and Exon-2 using specific primer pairs. We used Sanger sequencing to sequence the amplified fragments at the Exon-1 end. A highly accurate method with a 99.99% success rate. The sequences were then aligned with the NCBI GenBank database and analyzed using BioEdit 6.0 and MEGA 12.0 software. Our findings confirmed that the BMP15 gene is present in all the breeds we studied, but we didn’t find any polymorphisms among the Bangladeshi rams. The phylogenetic analysis showed strong genetic connections between local and international breeds. These results suggest the potential for enhancing Indigenous sheep through male-line genetic manipulation techniques, like selective semen use or artificial insemination, while preserving the BMP15 sequences to improve their productive traits.

## Introduction

The native sheep (Ovis aries) of Bangladesh originated from the wild Urial (Ovis Orientalis Vignei) of Asia. In the country, most sheep are of indigenous types, which are reared in the Barind, Jamuna basin, and Coastal areas (1). The traditional rearing system causes reduced meat production and poor reproductive performance, which causes economic losses for the sheep farmers (2). Despite the productive performance due to several technical (genotype, feeding, and health management), institutional, environmental and infrastructural constraints, Indigenous sheep breeds have great potential to contribute more to the smallholder who has low-input, crop-livestock and pastoral production systems (3).

Three genes associated to fertility were found in sheep: Growth distinction element 9 (GDF9), called FecG; bone morphogenetic protein receptor type 1B (BMPR1B), called FecB; and bone morphogenetic protein 15 (BMP15), known as FecX (4,5). Bone morphogenetic protein 15 (BMP15) is a member of the altering development factor-b (TGFB) superfamily that plays a necessary function in mammalian ovary improvement, oocyte maturation, and litter size. Bone morphogenetic protein 15 (BMP15/FecX) gene is thought about among the significant genes and a prospect marker for reproduction in animals, specifically sheep. BMPR1B was the very first gene recognized for its role in controlling prolificacy in sheep (6) and has mutation in some sheep breeds like Boorola Merino sheep (7) and Javanese sheep (8). Research studies have actually shown that mutation in BMP-15 can cause both increased ovulation rates and infertility (9). Associations in between mutations in the BMP-15 gene and reproductive performance have actually been extensively studied by many previous scientists (10,11). Recently, the expression and localization of BMP-15 in male testis and epididymis have been studied. Studies have actually revealed that BMP-15 is stage-specifically localized in X chromosome of gonads and have influence on pachytene stage of spermatocytes (12). The expression patterns of BMP-15 in the testis suggest that its proteins may have unique functions at defined points of spermatogenesis (13). On the other hand BMP-15 null mice are fertile and typical (14). BMP-15 must be considered as a potential hereditary marker for sperm quality, based on its association with fresh sperm motility (15).

To date, the expression of BMP15 mRNA in the testes has been determined in some mammalian species, including humans, rats, and mice (12,16,17) as well as in fish species such as the European eel and catfish (18,19). Recently, a connection was determined between BMP15 expression and the seasonal variation in testicular steroidogenesis (19). It has been demonstrated that the ovine gene BMP15 presents a mutation in the Rasa Aragonesa Spanish sheep breed, which has actually been called FecXR. This mutation produces high-quality semen and enhances some sperm specifications in this type, making these males specifically valuable for artificial insemination (20). Based on previous research studies, the biological function of BMP15 in male reproduction requires additional investigation.

Research studies on BMP15 in rams have examined tissue expression patterns in rams that vary in fecundity (21), fertility rate (22), and the impact of the FecB genotype on semen attributes (23). Little is known concerning the expression pattern and biological function of BMP15 in male gonads. The objective of this work was to identify the polymorphism of the BMP15 gene in rams from different sheep breeds of Bangladesh and find out the ancestor’s relationship with other worldwide sheep breeds and reproductive quality.

## Materials and Methods

### Animals and Blood Sampling

The research study work was conducted from January 2023 to December 2024 at the sheep farm of the Department of Surgery and Obstetrics, Bangladesh Agricultural University, and was approved by the Animal Welfare and Ethics Committee, following the animal principles rules (Approval No: AWEEC/BAU/2024(01), Dated 20.02.24).

The study involved twenty-five (25) rams (Table 1), which were collected from various areas across the country. These rams were raised for one year under a traditional system and were trained for semen production using the artificial vagina (AV) method, with a body condition score (BCS) of 3. All the animals were kept under the same nutritional conditions.

**Table 1:**
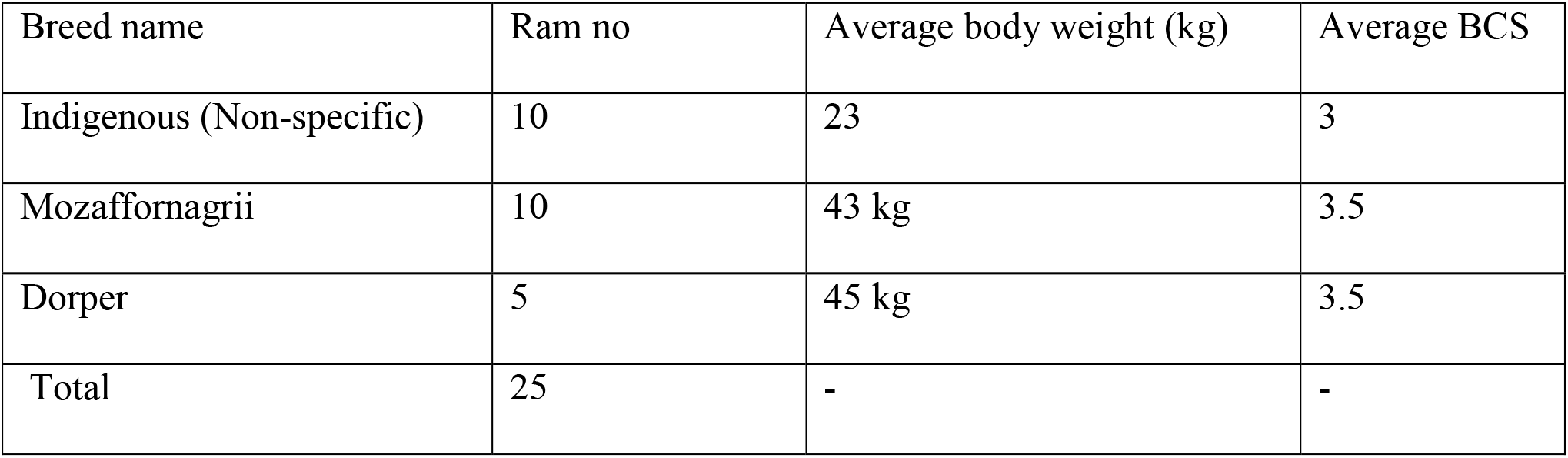
Rams description used for experiments.

Blood samples (5 mL) were collected from each animal through the jugular vein into EDTA-containing tubes. The blood samples were stored at − 20 ° C until DNA extraction. All samples were collected from the animals at the same time to guarantee consistency in the sampling procedure.

### DNA extraction and PCR amplification

Genomic DNA was extracted from whole blood utilizing the Reliaprep ™ Blood gDNA Miniprep System Kit, according to the Technical Manual of Promega Corporation. The DNA pellet was kept at − 20 ° C for further experiments. DNA concentration was measured using the NanoDrop approach (García-Alegría et al., 2020). Exon 1 and exon 2 of BMP15 were magnified using 2 set of specific primers, which were formerly developed by (24) and (15) (Table 2). The PCR amplifications were carried out in a thermal cycler (Gene Atlas, Model G2, Astec, Japan), using a specific annealing temperature level (Table 2).

**Table 2:**
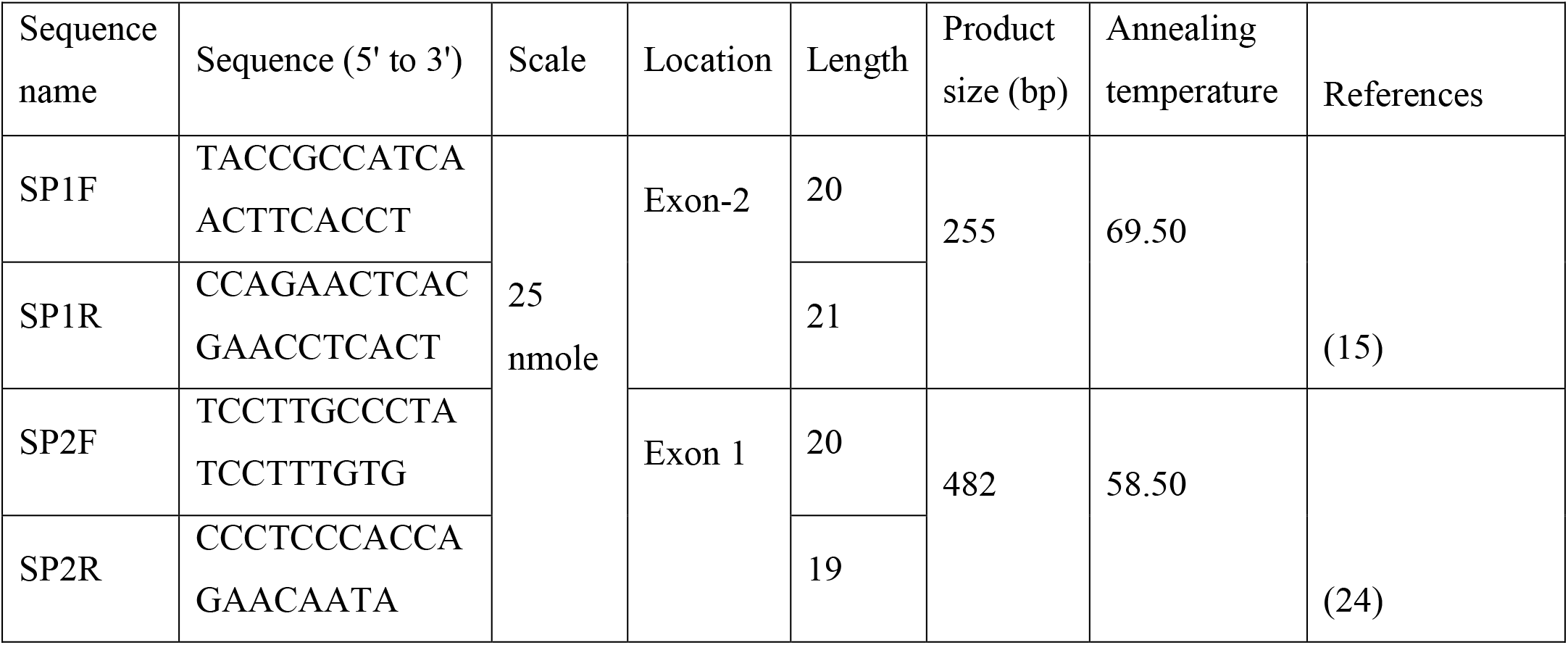
Primer and PCR condition utilized for BMP 15 gene detection and amplification.

### Sequencing of PCR Products

To find any possible polymorphism in exon-1, the PCR products from nine randomly chosen samples were sequenced. Consequently, PCR-RFLP (Polymerase Chain Reaction-Restriction Fragment Length Polymorphism) was used to confirm the mutations observed in the gene sequencing. After filtration of the PCR products, the samples were sent out to the Apical Scientific Laboratory in Selangor, Malaysia, for Sanger Sequencing. The sequencing was performed using the current automated genetic analyzer from Applied Biosystems ® following their enhanced protocol.

### Sequencing Analysis

The created raw series were examined utilizing direct sequencing with Bioedit software application (25). Series alignment was performed utilizing the NCBI/BLAST/blast suite. The series were aligned up separately with sequences from three different sheep breeds obtained from the National Center for Biotechnology Information (NCBI) utilizing ClustalW. The phylogenetic tree was then constructed based on the optimum possibility range utilizing MEGA 12.0 software application (26).

## Results

2 set of PCR products were resolved in 1% agarose gel electrophoresis for verification (band) separately at both Exon-1 and 2 of te BMP15 gene (Figure 1a and 1b). The PCR items were digested with AvaII limitation enzyme at 37 ° C for 5 h, and the resulting items were separated on a 2.5% agarose gel, pictured with ethidium bromide, and discovered by Alphaimager MINI gel paperwork system to visualize the image Figure 1a and 1b and the results were sown 100% positive indicating that all rams are fertile and prolific.

**Fig 1 a:**
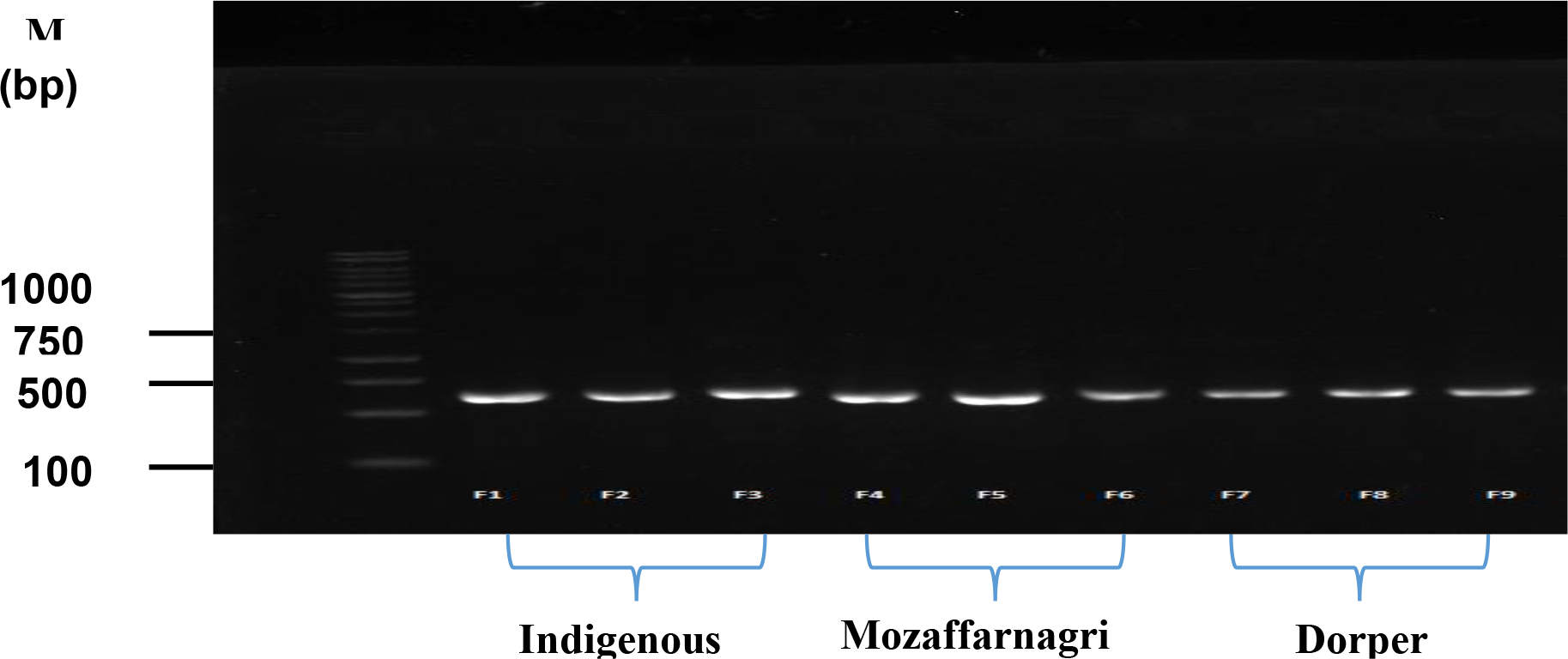
BMP15 detection at Exon-2 on 255 bp, M: denotes 100bp DNA ladder (Marker)

**Fig 1 b:**
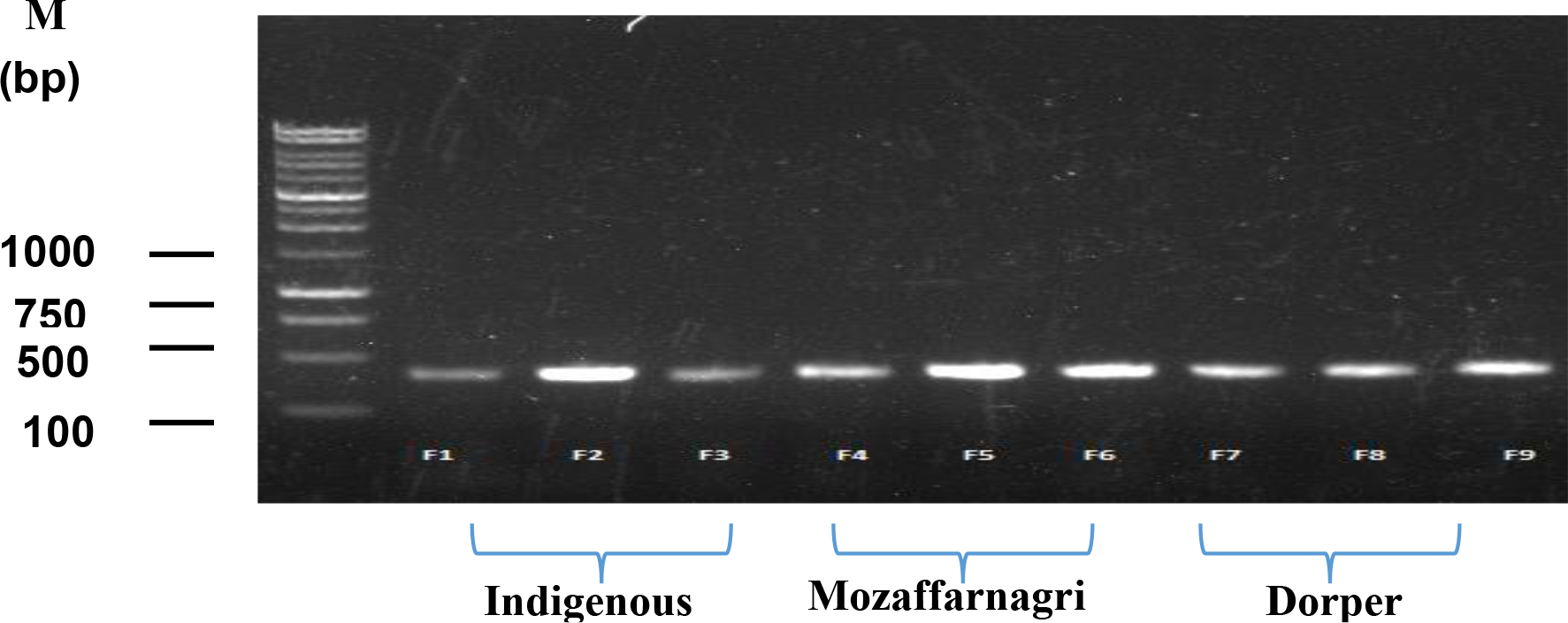
BMP15 detection at Exon-1 on 482 bp, M: denotes 100bp DNA ladder (Marker)

Exon 1 of the BMP15 Gene Sequence: From each of the 12 samples, exon 1 of the BMP15 gene was amplified and sequenced by an automated ABI DNA sequencer (design 373, PE Applied Biosystems). Comparison and blast information (Accession no: AY885263) showed that the E1 site had no polymorphism among the breed (2a, 2b, 2c). In Sanger Sequencing, outcomes were compared to NCBI/BLAST/blast suite positioning (accession no: AY885263), revealing 99% similarity (Figure 2b, 2a, and 2c). All types’ sequencing was similar. Anoter blast search of Exon-1 of BMP 15 DNA revealed a 100% match with Ovies aries (accession numbers AY885263) in the Sanger sequencing results sent to NCBI (Figures 2a, 2b, and 2c).

**Figure 2:**
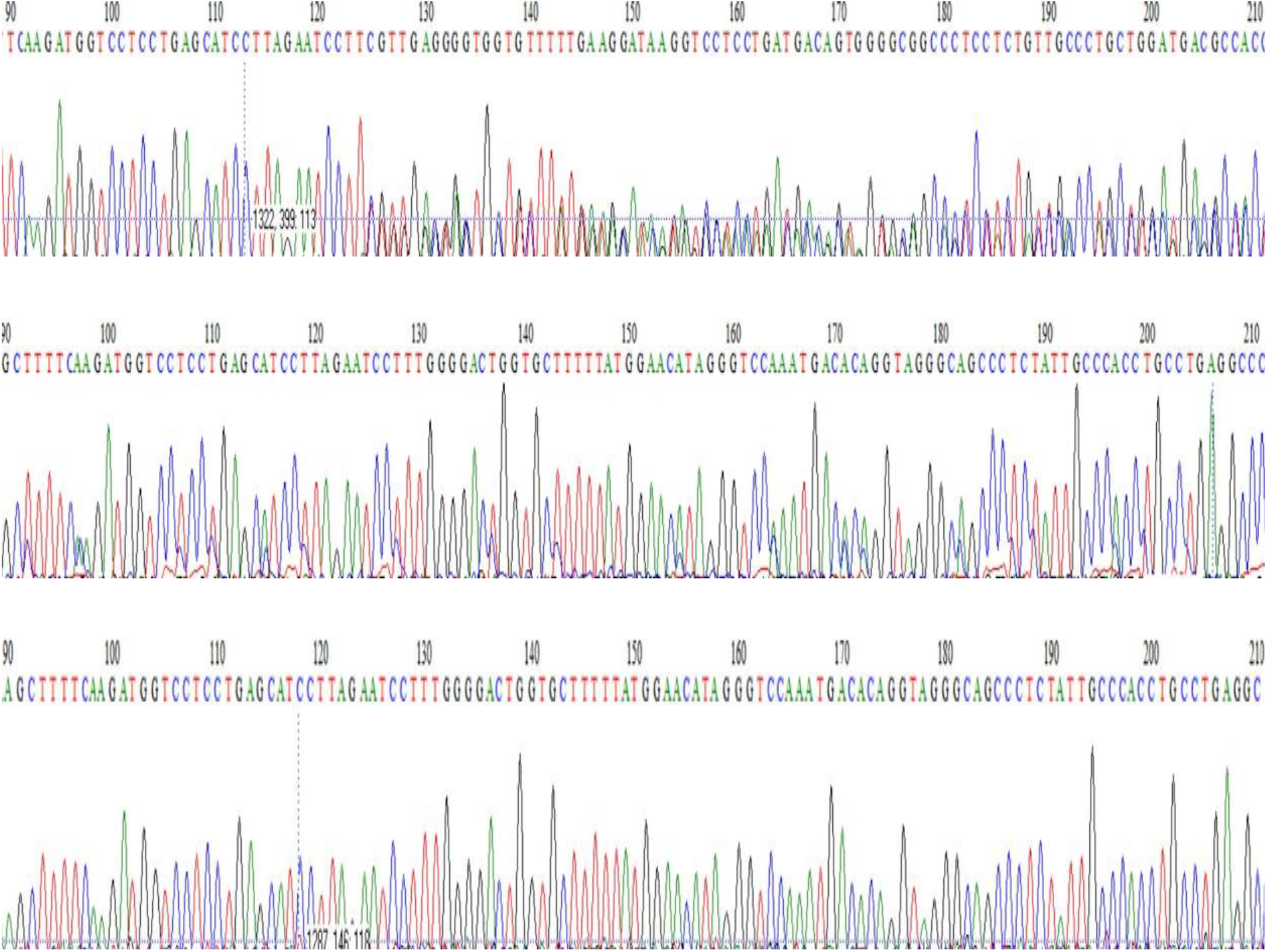
The chromatogram of BMP 15 gene at Exon-1 region sequencing of three different breeds a)Indigenous breed b) Mozaffonagri and c) Dorper breed

Evolutionary relationships of BMP15 gene: The phylogenetic analysis of BMP 15 were done from the haplotypes in this study. The phylogenetic tree for the study of evolutionary relationships of taxa of the BMP15 gene is shown in Figure 3. The outcome was only a 482-bp band, indicating that there was no polymorphism at this site in three ram breeds and are closely related to others world breed.

**Figure 3:**
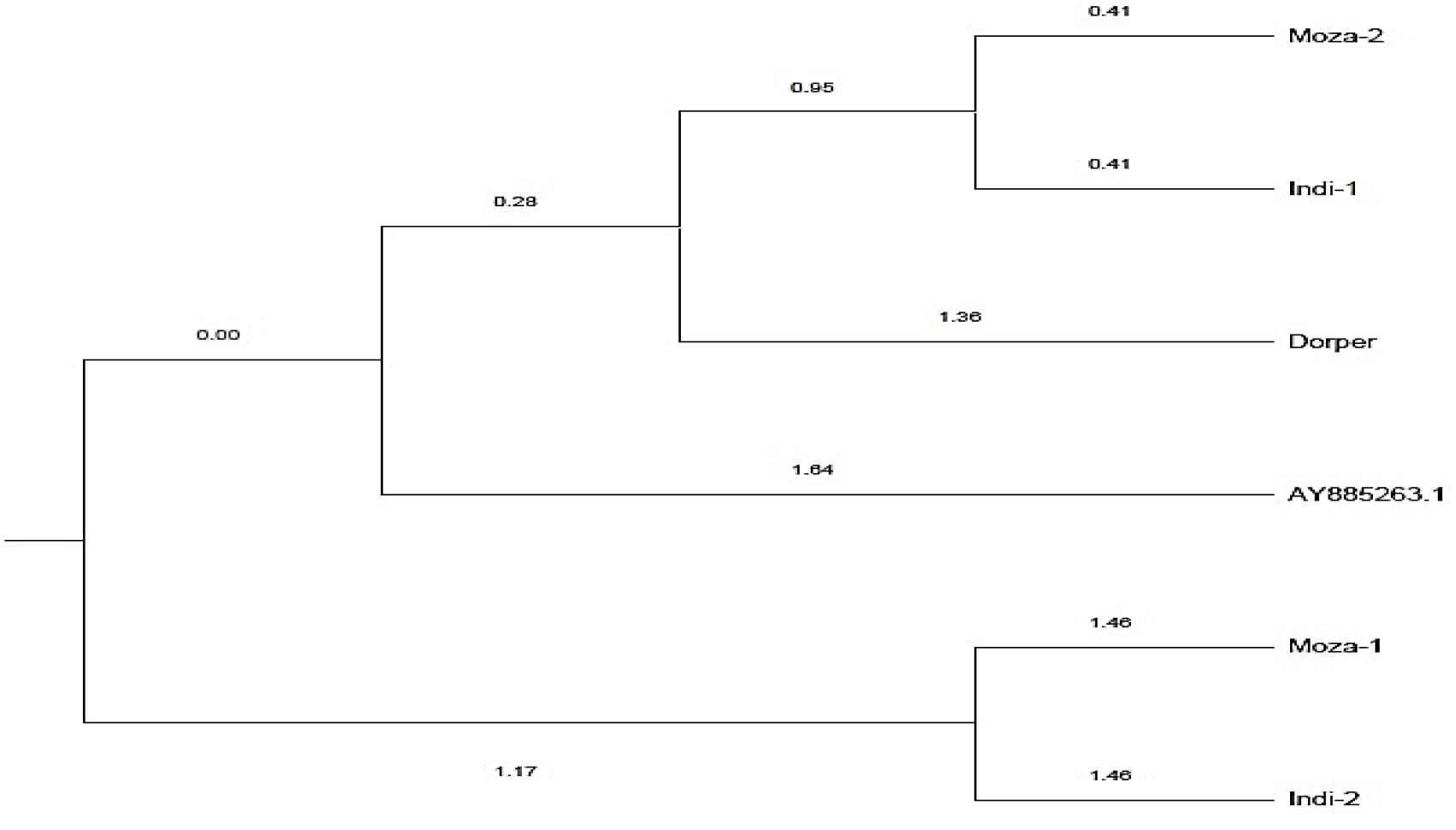
The UPGMA diagram of tree different breeds compared with a standard breed collected from NCBI blast.

## Discussion

Previous scientists have actually found that BMP-15 receptors are expressed by Sertoli cells and round spermatids, and GDF-9 can modulate key Sertoli cell functions (12). Sun et al., 2014 (15) suggested high levels of BMP-15 present in the bull testes might potentially regulate testicular function and has a putative association in between BMP-15 gene mutation and sperm quality traits in Chinese Holstein bulls.

Mutations of the BMP15 gene have been recognized in several sheep breeds across the world (27), showing that the heterozygous genotype (B+) affects the additive effect on ovulation rate.

Dube et al., 1998 (28) found mutations in the BMP15 gene, has result in high yields of Inverdale sheep. In some research studies, the G4 and G6 polymorphisms determined through sequencing agreed with those reported in sheep of the Belclare, Cambridge, Mehraban (5,29) and Barki, Osseimi, Rahmani, Saudanez, and Awassi breeds (30). In this study no mutations were discovered in the Exon-1 codifying regions of BMP15, according to the sequence analysis of this research study compared to those of the referral (accession no: AH009593.2 and KT238844.1). This statement is opposite to the above research results but supported by Khan et al., 2022 (31) who found the absence of polymorphism in ewes on the Exon-2 region.

### Phylogenetic analysis

To discover the evolutionary relationship of Bangladeshi indigenous sheep with other sheep breeds or populations, 2 different approaches were employed for phylogenetic tree building. The very first technique was based upon reconstructed haplotypes of indigenous sheep of Bangladesh in addition to Ovis aries as a referral (Figure 3). The second technique considered haplotype information from Bangladeshi native sheep, different sheep types or populations all over the world, and other sheep types (Figure 4).

Phylogenetic analysis exposed close relationships amongst sheep from different countries without forming particular clusters or groups for any particular population. More specifically, one significant cluster consisted of almost 60% of Bangladeshi sheep population data from all considered regions along with Ovis aries reference information, determining indigenous sheep as the sole progenitors of Ovis aries (Figure 3) also found in Mousumee et al., 2021 (32) research. In India Garole and Nagpuri sheep formed different clusters without significant distinctions (33,34). These findings are supported by the present study.

The absence of any distinct hereditary structure amongst the studied populations recommended strong gene flow due to intermixing amongst Bangladeshi sheep populations or common maternal family trees among the regions (35). Figure 3 reveals that native sheep of Bangladesh formed a cluster exclusively with Ovis aries species, illustrating their single ancestor.

More especially, the phylogenetic analysis showed close relationships in between a lot of Bangladeshi sheep samples and sheep types from India (Garole and Bonpala), China (Tibetan), and due to quickly spread out from their domestication centre in Southwestern Asia following human migrations (36).

Finally, these findings were constant with the phylogeny outcomes of a previous research study on the BMP15 gene by Bibinu et al. (2016)(37), which reported comparable clustering of sequences amongst various types in spite of some intermingling in between species. The phylogenetic relationship showed that indigenous Bangladeshi sheep resemble sheep from Mexico, North America, and Egypt. This resemblance may be credited to shared ecological conditions and ancestral relatedness, showing evolutionary processes in these sheep.

## Conclusion

The genotypic diversity of sheep breeds in Bangladesh was discovered to be low based on a series variation analysis. According to this study, low genetic distinction in among the studied three sheep breeds can be credited to a shared maternal lineage and significant gene flow triggered by interbreeding. An analysis of the phylogenetic tree of Bangladeshi indigenous sheep exposed that they descend from a single forefather, Ovis aries. Bangladesh’s indigenous sheep are the subject of the very first study on ram BMP15 results, which can be utilized to improve conservation and reproducing initiatives in the future through.

## Notes

### Competing Interest Statement

Yes, I will do after accepatnce.

